# TDP 2 modulates the expression of estrogen-responsive oncogenes

**DOI:** 10.1101/2022.06.01.494417

**Authors:** Nicholas Manguso, Minhyung Kim, Neeraj Joshi, Rasel Al Mahmud, Juan Aldaco, Ryusuke Suzuki, Felipe Cortes-Ledesma, Xiaojiang Cui, Shintaro Yamada, Shunichi Takeda, Armando Giuliano, Sungyong You, Hisashi Tanaka

**Affiliations:** Department of Surgery, Cedars-Sinai Medical Center, West Hollywood, CA, 90048, USA; Department of Radiation Genetics, Graduate School of Medicine, Kyoto University, 606-8501 Kyoto, Japan; Centro Andaluz de Biologia Moleculary Medicina Regenerativa (CAMBIER), CSIC-Universidad de Sevilla (Departmento de genetica) Sevilla, Spain; Biomedical Sciences, and Cedars-Sinai Medical Center, West Hollywood, CA, 90048, USA; Cedars-Sinai Cancer, Cedars-Sinai Medical Center, West Hollywood, CA, 90048, USA

## Abstract

With its ligand estrogen, the estrogen receptor (ER) stimulates tumor cell growth by activating a global transcriptional program. This activation involves topoisomerase 2 (TOP2), which resolves topological problems by transiently creating and re-ligating DNA double-strand breaks (DSBs). Recent studies have uncovered the involvement of DNA repair proteins in the repair of TOP2-induced DSBs. These noncanonical repair pathways may serve as backup processes when TOP2 is halted and fails to re-ligate DSBs, but their impact on transcription remains elusive. In this study, we investigated the role of tyrosyl-DNA phosphodiesterase 2 (TDP2), an enzyme that acts for the removal of halted TOP2 from the 5’-end of the DNA, in the estrogen-induced transcriptome. Using TDP2-deficient ER-positive cells and mice, we showed that TDP2 regulates the expression of oncogene *MYC*. *MYC* induction by estrogen was a very early event (1 hour) and TOP2-dependent. In TDP2-deficient cells, the induction of *MYC* by estrogen became prolonged and volatile. Bulk and single-cell RNA-seq identified the oncogenes *MYC* and *CCND1* as genes whose estrogen response was regulated by TDP2. These results suggest that TDP2 may play a role in the repair of TOP2-induced DSBs in specific genomic loci and tightly regulates the expression of oncogenes.

## Introduction

With a lifetime risk of one in eight women, breast cancer stands as the most prevalent female cancer, remaining the most frequent cause of cancer-related death in women. Approximately two-thirds of breast tumors express estrogen receptor alpha (ERα), and their growth depends on the hormone estrogen [1]. Estrogen plays a crucial role in the development and regulation of female reproductive systems and secondary sex characteristics. 17beta-estradiol (E2) is the predominant estrogen during reproductive years both in terms of absolute serum levels and activity. As with all steroid hormones, secreted E2 diffuses into the cell nucleus, where it binds to a receptor ERα, forming E2-ERα complex [2]. The E2-ERα complex subsequently binds to DNA, acting as potent regulators of gene expression. Transcriptional activation *in vitro* occurs within an hour after E2 stimulation and involves Type II topoisomerase (TOP2), specifically, TOP2 beta (TOP2β) [3, 4].

TOP2 activities are crucial in managing torsional stress during various processes of DNA metabolism, including replication, chromosome segregation, and transcription [5, 6]. It catalyzes to induce a transient DNA double-stranded break (DSB) in a DNA duplex, with passing another intact duplex through the broken DNA. This process involves a transient intermediate with a covalent bond between the catalytic tyrosine in TOP2 and the 5’ phosphate groups of the DNA ends (TOP2 cleavage complex, TOP2cc), followed by the re-ligation of the DSB ends. In the context of general transcription, advancing RNA polymerase II (PolII) on the DNA templates creates over-wound DNA in front and leaves under-wound DNA behind. By relieving torsional stress in front, TOP2 promotes the release of paused PolII for transcriptional elongation [7–9]. Because of the important and broad roles of TOP2 in nucleotide metabolisms, small molecule inhibitors that intervene in each step of TOP2 activity have been developed. TOP2 catalytic inhibitors prevent the induction of DSBs, while TOP2-poisons, by stabilizing TOP2cc, prevent the re-ligation of TOP2-induced DSBs [10].

In addition to relieving torsional stress during general transcription, TOP2 plays a crucial role in the induction of a specific set of genes in response to exogenous stimuli, including E2, heat shock, serum starvation, and neuronal activity [7, 9, 11, 12]. On some of these gene loci, TOP2β was shown to induce DSBs at transcriptional units, including promoters, in response to these stimuli, and following the DSBs were the recruitment of DSB repair proteins, DNAPK and Ku70 [9, 11]. This recruitment is intriguing, considering that TOP2 is established as a self-sufficient enzyme, and DSBs induced by TOP2 are transient and usually re-ligated by TOP2 [5, 13]. TOP2β occupies over 2,000 genomic sites upon E2 treatment in MCF7 breast cancer cells [14], suggesting a widespread involvement in transcriptional regulation. A fraction of these sites would possibly recruit DSB repair proteins, as noncanonical repair processes for Top2-induced DSBs. Two classes of repair proteins have been identified; (1) proteins involved in processing (removing) TOP2cc that remains covalently attached to the 5’ DSB ends and (2) those repairing the resulting clean DSB ends. The first process was executed either by 5’-tyrosyl-DNA phosphodiesterase (TDP2) or the Mre11-RAD50-Nbs1 (MRN) complex [15–17]. The resulting clean ends are then repaired by non-homologous end-joining (NHEJ) [18]. BRCA1 is also implicated in the repair of DSBs with TOP2cc [17, 19]. Although these noncanonical processes are currently viewed as backup processes, for instance where TOP2 is trapped at the DNA ends and requires assistance, it remains elusive whether noncanonical processes are inherently programmed to contribute to the repair of TOP2-induced DSBs in specific gene loci.

Intriguingly, when left unrepaired, TOP2-induced DSBs can activate a specific set of genes. For example, inhibiting the re-ligation of TOP2-induced DSBs by TOP2-poison etoposide resulted in the upregulation of early response genes to neuronal activity in cultured primary neurons [11]. Understanding the mechanism behind the upregulation is crucial. It is also essential to identify which genes are upregulated in response to external stimuli when the repair of TOP2-induced DSBs is hindered. The upregulation of target genes can pose a serious problem, particularly when external stimuli, such as E2, promote cell proliferation. Such activation could potentially lead to abnormal cell proliferation and, ultimately, contribute to tumorigenesis. However, it remains unknown whether such upregulation can occur in a more natural setting of DSB repair deficiency, for example, in cells lacking a DSB repair protein, compared to etoposide treatment. Stalled TOP2cc by etoposide occurs very frequently in the entire cell population, enabling the detection of transcriptional upregulation using techniques for measuring the average gene expression levels in a cell population, such as qPCR, digital PCR, and bulk RNA-seq. In contrast, in cells deficient in a DSB repair protein, stalled TOP2cc may not occur throughout the entire cell population. In such a case, the aforementioned techniques might not detect transcriptional upregulation. However, upregulation could still be observed in DSB repair-deficient cells if TOP2-induced DSB is programmed to involve the DSB repair protein at a certain gene locus. This possibility was shown at the protein level in mice lacking NHEJ repair; E2 stimulation induced MYC protein in mammary epithelial cells of mice lacking DNA-PKcs [20].

We aimed to identify a set of E2-induced genes that become upregulated in cells deficient in TOP2-induced DSB repair. This gene set could also inform a potentially novel process by elucidating gene loci where noncanonical repair of TOP2-induced DSBs is programmed. To achieve this aim, we established ER-positive breast cancer cell lines lacking TDP2 and investigated their global transcriptional program. We chose TDP2 because TDP2 knockout cells/mice are viable [18] and amenable to E2 stimulation experiments. Using digital PCR and bulk- and single-cell RNA-seq, we identified that *MYC* was among several genes that exhibited enhanced transcription in response to E2 in TDP2-deficient cells compared to TDP2-proficient cells. We also found that another breast cancer oncogene, *CCND1*, showed increased transcription in TDP2-deficient cells. These results suggest that noncanonical repair processes of TOP2-induced DSBs may tightly regulate oncogenic gene expression. The results may also explain why deficiency of DSB repair proteins is often associated with the susceptibility of breast tumors.

## Results

### *MYC* induction by 17beta-estradiol (E2) depends on TOP2

*MYC* is a well-studied oncogene and known to transcriptionally respond to E2 [21]. To assess this response, we collected RNA from two ER-positive breast cancer cell lines MCF7 and T47D, before and after treatment with E2 (**Fig. 1A**). We employed digital PCR assays to measure the expression of *MYC* mRNA. In this assay, a *MYC* Taqman probe labeled with a dye FAM and a control probe, either *TBP* or *TFRC,* labeled with a dye VIC were mixed with cDNA from cell lines and PCR-amplified in approximately 20,000 x 1 nL reaction (**Fig. 1B**). *MYC* expression levels ( the number of *MYC*-positive 1 nL reactions) were normalized by the internal control (the number of internal control-positive reactions). We added 10 nM E2 and incubated cells for two hours before extracting RNA. The fold induction of *MYC* was calculated as the ratio of normalized *MYC* expression with E2 treatment to normalized *MYC* expression without E2 treatment.

**Figure 1.**
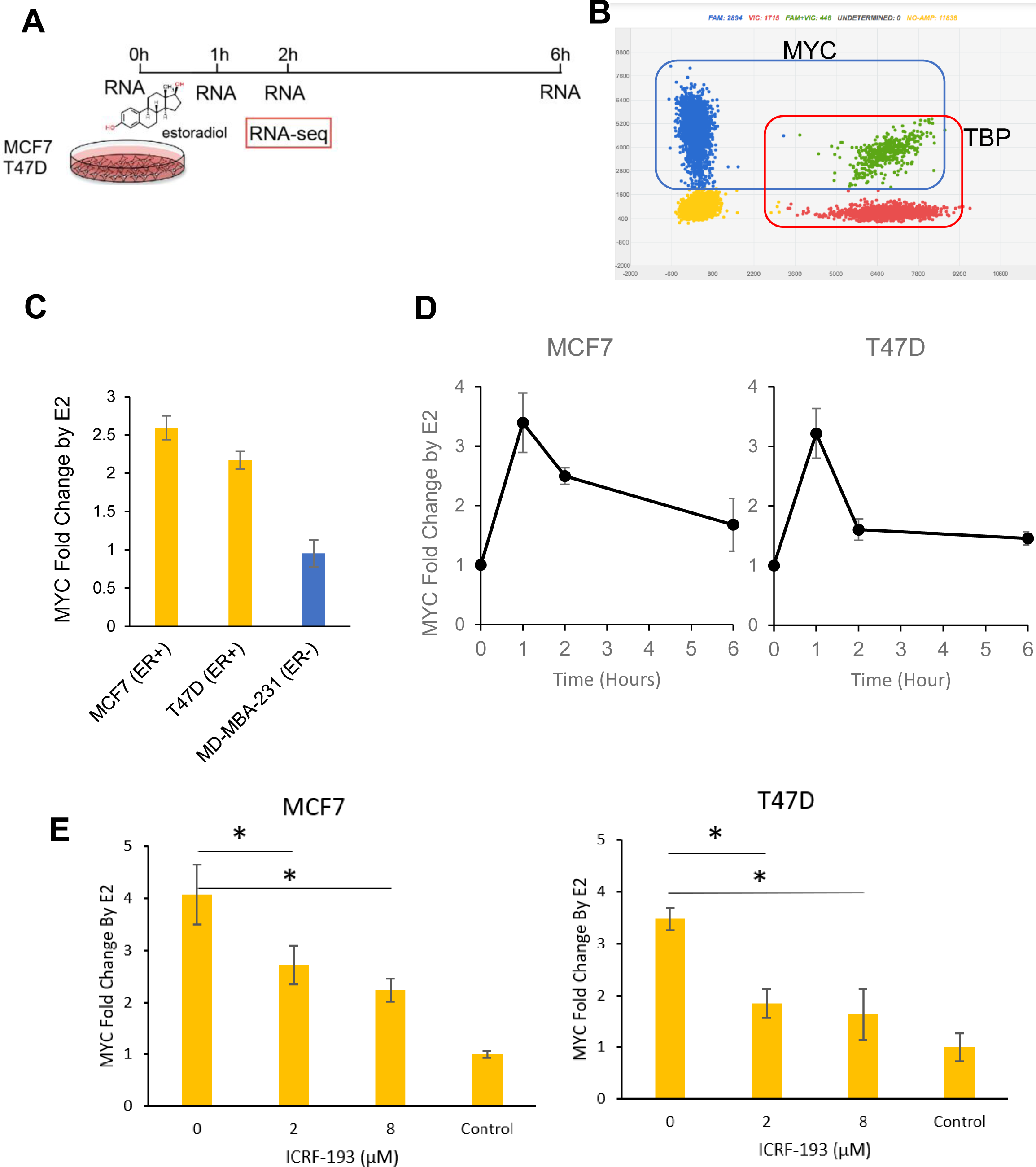
Estrogen and TOP2-dependent induction of *MYC*. (A) Experiment design: MCF7 and T47D cells were treated with 10 nM 17β-estradiol for 1, 2, and 6 hours. RNA was extracted at each time point for 3D PCR analysis, and bulk- and single-cell RNA-seq was performed at the 2-hour time point. (B) 3D PCR output from Analysis Suites. Blue dots represent wells only with amplification of genes detected by FAM (*MYC*) probe, red dots represent wells only with the amplification detected by VIC (*TBP* or *TFRC*) probe (internal control), yellow dots represent wells without amplification, and green dots represent wells with the amplification of both probes. Normalized *MYC* expression was calculated by the ratio of FAM (*MYC*) to VIC (*TBP* or *TFRC*). (C) *MYC* induction in cells after 2 hours of treatment with 10 nM E2 relative to cells with no E2 in two ER-positive breast cancer cell lines (MCF7 and T47D) and an ER-negative cell line (MD-MBA-231). Averages of three independent experiments are shown. (D) Dynamic responses of *MYC* induction: Cells were treated with 10 nM E2 for 0, 1, 2, and 6 hours. Averages of two independent experiments are shown. (E) A TOP2 catalytic inhibitor ICRF-193 Effects on *MYC* mRNA induction: MCF7 and T47D cells were treated with 2 and 8 μM ICRF-193 for 1 hour, followed by the treatment with 10 nM E2 for 2 hours. *MYC* induction decreased by ICRF-193 in both cell lines. Averages of three independent experiments are shown. * p<0.05, ** p<0.01, Student’s t-test.

ER-positive (ER+) cell lines, MCF7 and T47D, showed an increase in *MYC* expression after E2 treatment (2.59-fold and 2.17-fold, respectively). As expected, the ER-negative (ER-) cell line MDA-MB-231 did not show the induction (0.93-fold change) (**Fig. 1C**). To understand the temporal regulation of *MYC* expression, we measured *MYC* mRNA at 1, 2, and 6 hours after the addition of 10 nM E2. In both the MCF7 and T47D cell lines, *MYC* expression peaked at 1 hour (MCF7 2.89-fold, T47D 2.79-fold), followed by gradual decreases until 6 hours (**Fig. 1D**). As previous studies reported that TOP2 mediates the induction of E2-responsive genes [3], we then tested whether *MYC* induction is also TOP2-dependent. We used a TOP2 catalytic inhibitor, ICRF-193, which prevents TOP2 from creating DSBs [22, 23]. We added ICRF-193 to the cell culture media 1 hour before adding 10 nM E2 and incubated cells for 2 hours after E2 treatment. We observed a dose-dependent inhibition of *MYC* induction in both MCF7 and T47D (Figure 1E), strongly suggesting that E2 stimulation of *MYC* expression in ER-positive cells is TOP2-dependent.

### *MYC* transcriptional activation in TDP2 knockout cells

Since ICRF-193 inhibits DSB induction, DSBs may play a pivotal role in the transcriptional activation of *MYC*. When left unrepaired, TOP2-induced DSBs promote transcription for a specific set of genes [11]. Therefore, our next objective was to determine whether unrepaired DSBs could further enhance *MYC* induction by E2. To test the idea, we knocked out *TDP2* in ER-positive cell lines. When TOP2 fails to re-ligate DSBs and is aborted, TOP2cc is processed by a ubiquitin-proteasome pathway or, alternatively, a proteosome-independent pathway [24, 25]. TDP2 removes the remnant aborted TOP2 and creates a clean DSB for NHEJ machinery to repair [18]. We used CRISPR-Cas9 to target *TDP2* in both MCF7 (MCF7 TDP2KO-1) and T47D cell lines. Additionally, we employed an independently created TDP2 knockout MCF7 cell line (MCF7 TDP2KO-2)[17]. We confirmed the lack of TDP2 protein in knockout cells by western blotting (**Fig. 2A**). The functional TDP2 deficiency was also confirmed using the TOP2 poison, etoposide. Etoposide traps TOP2cc at DSBs, which would require TDP2 to repair. Both the MCF7 and T47D cells deficient in TDP2 were significantly more sensitive to etoposide than parental cell lines (**Fig. 2B**).

**Figure 2.**
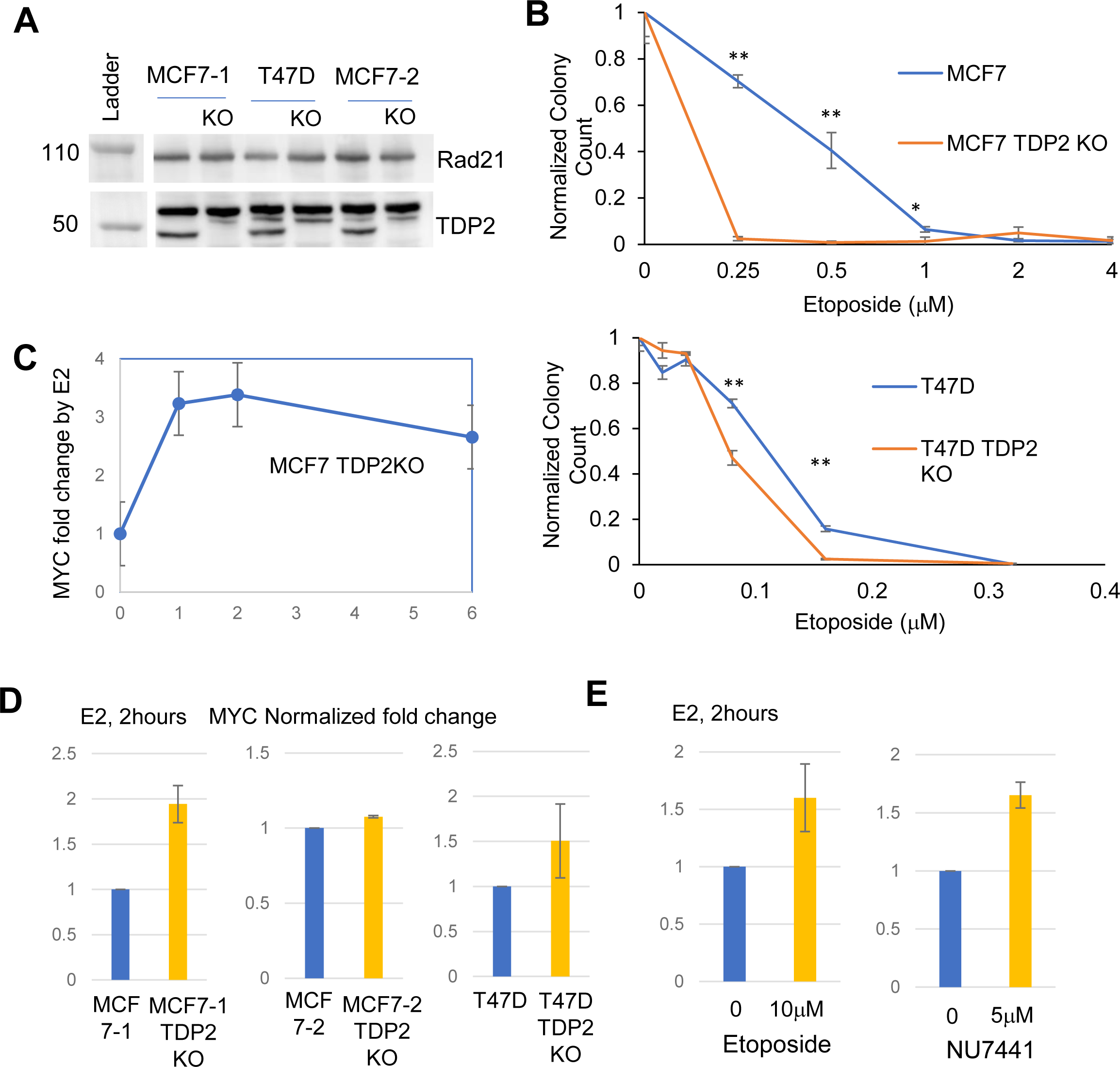
TDP2 suppresses the volatile and prolonged induction of *MYC* by estrogen. (A) Western Blot Analysis displaying MCF7-1, T47D and MCF7-2 cell lines alongside TDP2 KO cells for each cell line. (B) Etoposide sensitivity measured by colony formation assay for MCF7 and MCF7 TDP2KO(top) and T47D and T47D TDP2KO (bottom). Results from three independent experiments. * p<0.05, ** p<0.01, Student’s t-test (C) Persistent *MYC* transcriptional responses: MCF7 and MCF7 TDP2 KO cells were treated with 10 nM E2 for 1, 2, and 6 hours. Averages from two independent experiments. (D) *MYC* transcriptional responses at 2 hours: MCF7-1, MCF7-2 and T47D, and their TDP2 KO counterparts were treated with 10 nM E2 for 2 hours, and RNA was extracted for digital PCR assays. Results from two independent experiments. (E) NU7441 and etoposide increased the *MYC* transcriptional response. MCF7 cells, cultured with or without NU7441 or etoposide for three hours were then treated with 10 nM E2 for 2 hours. Results from two independent experiments.

We monitored the dynamics of *MYC* induction using digital PCR, as illustrated in **Figure 1C**. The highest level of induction at 1 hour after the addition of E2 (10nM) was similar between MCF7 and MCF7 TDP2KO-1 cells (**Fig. 2C**). However, *MYC* expression continued to be high in MCF7 TDP2KO cells, whereas it declined quickly in MCF7 cells. We further measured the induction of *MYC* at 2 hours after the addition of 10 nM E2 and found the increased fold induction of *MYC* in all three TDP2 knockout cell lines relative to their parental TDP2 wild-type counterparts (**Fig. 2D**). Because the expression was measured by total RNA of entire cell populations, these results suggest that *MYC* induction by E2 was enhanced in a significant proportion of the TDP2KO cell population. The prevailing view is that alternative repair proteins, including TDP2, only repair abortive TOP2-induced DSBs that could occur in a very small fraction of cells. However, these results suggest that a significant fraction of cells require TDP2 to regulate *MYC* induction. The delayed repair of TOP2-induced DSBs could promote the *MYC* response to E2 at 2 hours, as previously described for early response genes to neuronal activity [11]. Consistently, we observed that etoposide promotes the induction of *MYC* in MCF7 cells (**Fig. 2E**, left). Inhibiting NHEJ could also promote *MYC* response, as previously shown *in vivo* in DNA-PKcs deficient mice [20]. To explore this possibility, we inhibited NHEJ by DNA-PKcs inhibitor NU7441 [26]. Indeed, *MYC* induction at 2 hours after the addition of E2 was increased by NU7441 (5 μM) (**Fig. 2E**, right).

### Induction of MYC by estrogen *in vivo* in mice mammary glands

To assess the relevance of the above observation *in vivo*, we examined *MYC* response to E2 at the protein level in mice mammary glands. We administered E2 (300 μg/kg body weight) or PBS (control) intraperitoneally into three *Tdp2* wild-type and *Tdp2^-/-^* mice [18]. Mice were examined before injection (0 hours) and 6 and 20 hours after injection for MYC protein expression in CK8/18-positive mammary epithelial cells by immunostaining (**Fig. 3A**). We quantified MYC-positive epithelial cells in 3 to 12 mammary ducts at each timepoint and scored the fraction of MYC-positive cells in each mammary duct (**Fig. 3B**). MYC-positive cells increased in both wild-type and *Tdp2^-/-^* mice at 6 hours. MYC-positive cells become less frequent at 20 hours, showing transient induction of MYC in mammary epithelial cells. MYC-positive cells in *Tdp2^-/-^*mice at 20 hours were significantly more abundant than in *Tdp2* wild-type mice (p<0.01) at 20 hours or in *Tdp2^-/-^* mice without E2 at 0 hours (p<0.01). The prolonged induction of MYC in *Tdp2^-/-^* mice parallels the observation in transcription in TDP2KO MCF7 cells (**Fig. 2C**) and aligned with our *in vitro* findings.

**Figure 3.**
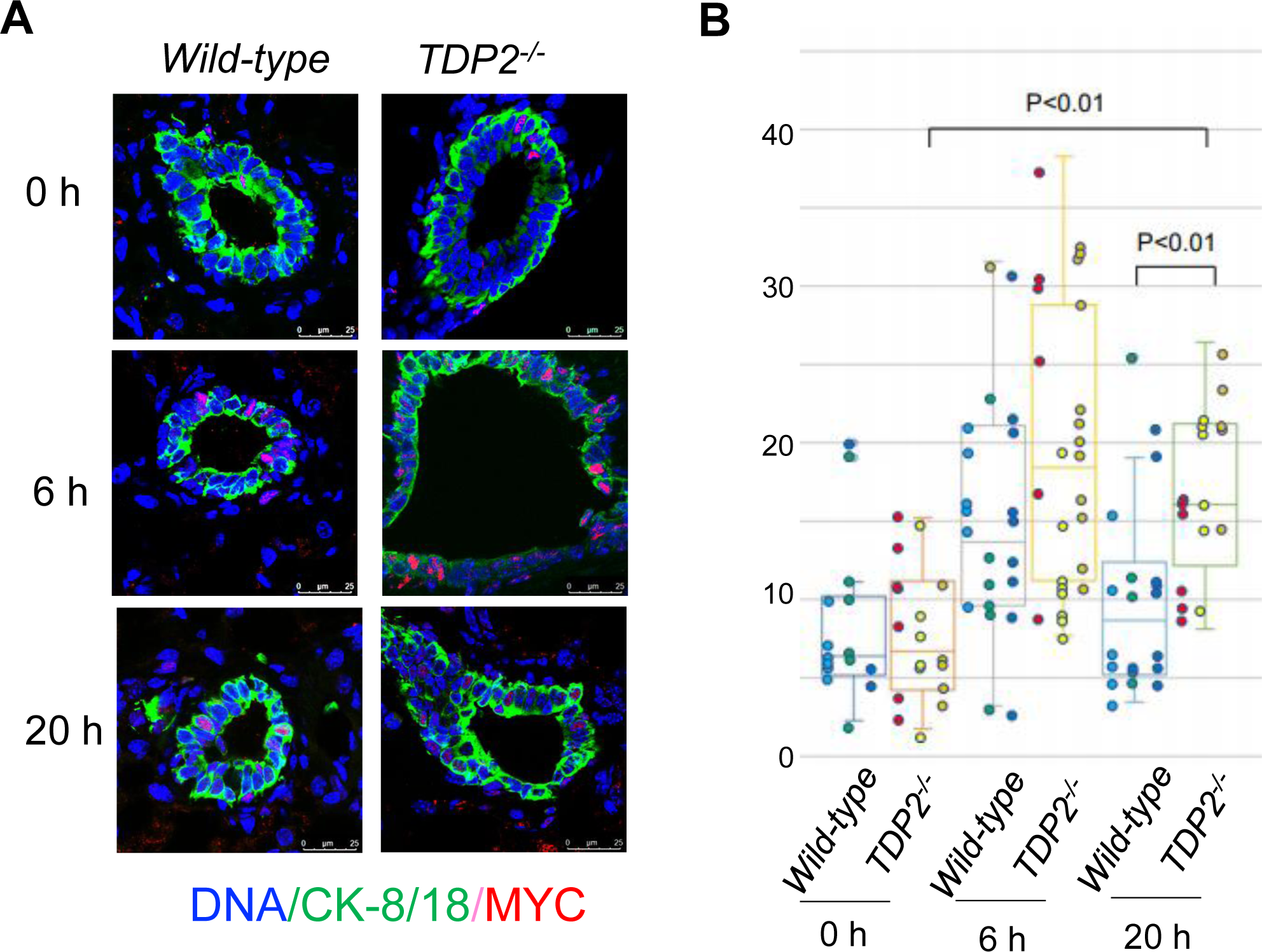
The induction of MYC protein by E2 *in vivo*. (A) Representative images of mammary ducts: Shown are images of mammary ducts from wild-type (left) and TDP2KO (right) mice, immunostained for MYC (red) and CK-8/18 (green) proteins. The images were captured before injection (0 hours) and 6 and 20 hours after the injection of E2. (B) Box plots illustrating the fraction of MYC-positive cells: Each plot represents Each plot represents the distribution of the fraction of MYC-positive cells among CK8/18-positive mammary epithelial cells. MYC-positive epithelial cells were scored in 3 to 12 mammary ducts in each mouse at each timepoint. p values were determined by t-test.

### MYC and MYC downstream targets induced by E2

MYC is a beta Helix-Loop-Helix transcription factor that, in conjunction with a partner protein Max, binds to DNA at the CACGTG E-Box sequence, thereby modulating global gene expression [27]. To examine the expression of MYC-regulated genes in MCF7 and MCF7 TDP2KO cells, we conducted RNA-seq analysis using total RNA collected before and 2 hours after the addition of 10nM E2 (bulk RNA-seq). We found a large number of differentially expressed genes in both MCF7 (2,021 genes) and MCF7 TDP2 KO cells (1,034 genes) (**Tables S1** and **S2**; fold change ≥ 1.5 and combined P-value < 0.05). Notably, *MYC* showed upregulation in both cell types, with a more pronounced increase in MCF7 TDP2KO cells (3.16 fold) compared to parental MCF7 cells (1.94 fold) by E2 stimulation (**Fig. 4A**), validating the prolonged response of *MYC* to estrogen in the absence of TDP2. *MYC* transcriptional response was slightly more pronounced in MCF7 cells treated with etoposide (2.17 fold, **Table S3**), suggesting the unrepaired TOP2-induced DSBs contributed to the enhanced response.

**Figure 4.**
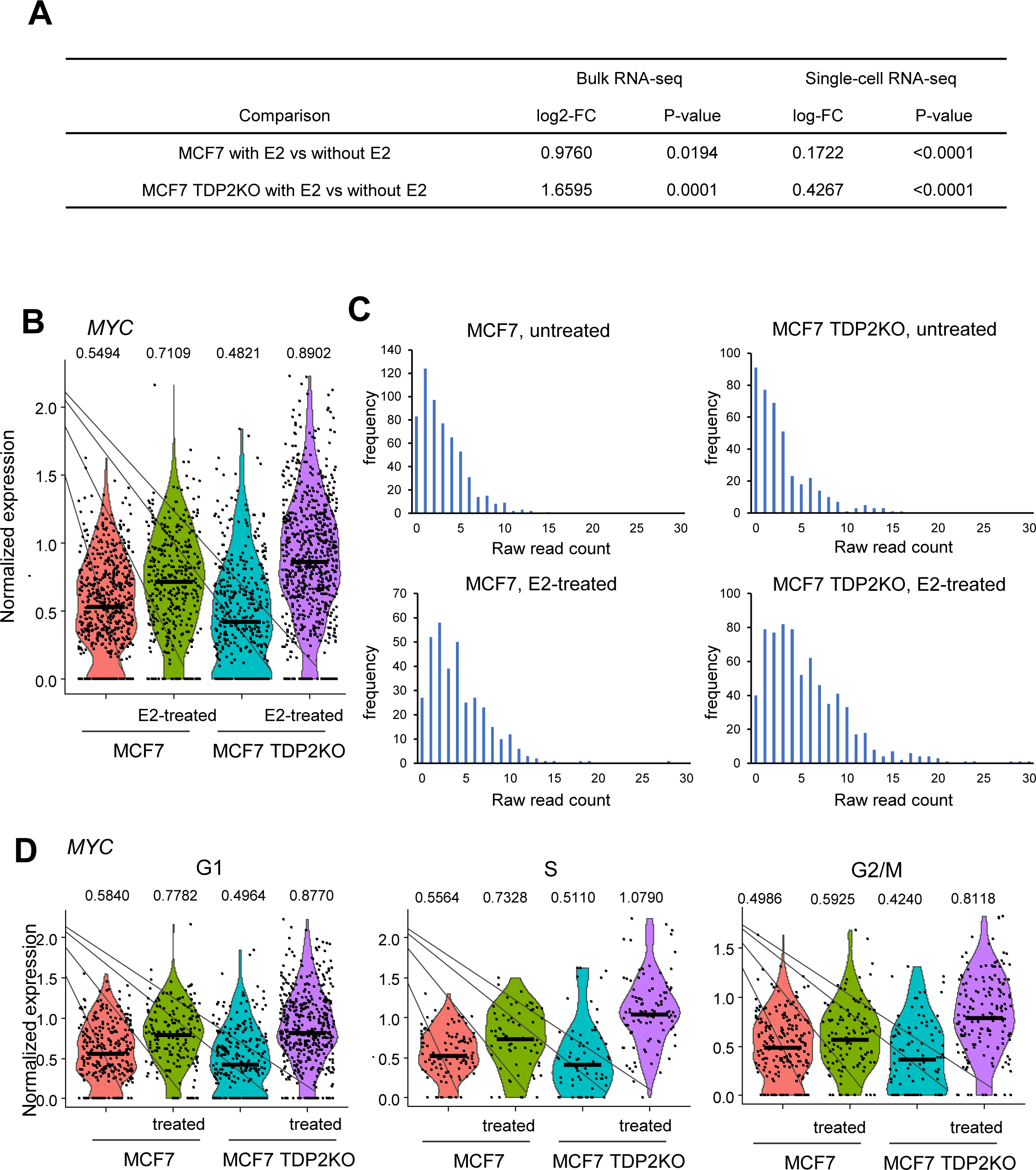
Bulk and Single-cell RNA sequencing analysis of *MYC* transcriptional response by E2. (A) *MYC* transcriptional response to E2 (2 hours) in MCF7 and MCF7 TDP2KO cells, measured by bulk RNA-seq and single-cell RNA-seq. (B) Violin plots displaying *MYC* expression: Normalized *MYC* expression distributions in MCF7 and MCF7 TDP2KO cells with or without E2 (2 hours) from single-cell RNA-seq data. (C) Histograms illustrating *MYC* expression distribution: Showing the skewed unimodal distribution of MYC expression in MCF7 and MCF7 TDP2KO cells with or without E2 (2 hours), obtained from single-cell RNA-seq data. (D) Violin plots of *MYC* expression at single-cell level: Depicting the normalized expression of *MYC* in G1, S, and G2/M cell populations.

To assess the outcomes of *MYC* activation following E2 treatment in MCF7 TDP2KO and parental MCF7 cells, we performed Gene Set Enrichment Analysis (GSEA) [28] with predefined MYC target genes. GSEA determines whether a defined set of genes shows statistically significant, concordant differences between two biological states. Specifically, we examined the significant changes in the expression of genes whose promoters harbor MYC binding sites within the 4 kb of the transcription start sites (MYC_Q2), potentially representing direct targets of MYC (**Fig. S1, A**, top). Upon treating cells with E2 for 2 hours, the majority of genes in MYC_Q2 exhibited nonrandom distribution, predominantly showing upregulation in both MCF7 and MCF7 TDP2KO cells. We determined the normalized enrichment score (NES) (**Fig. S1, B**), which reports the degree of overrepresentation of genes at the extremes of all the ranked genes. We found that MYC_Q2 genes were slightly more clustered in highly upregulated genes in MCF7 TDP2KO cells than in MCF7 cells. Furthermore, we examined gene sets whose expression was enhanced by overexpressing MYC in human primary mammary epithelial cells (Hallmark MYC target v2 and MYC_UP.V1_UP, **Fig. S1A** and **B**) [29]. Both sets demonstrated he enrichment of MYC targets in upregulated genes, with comparable NES values between MCF7 and MCF7 TDP2KO. These results suggest that the upregulation of MYC by E2 modulated global transcriptional programs in both cell types.

### MYC induction at the single-cell level

Bulk RNA-seq offers insights into the average RNA expression across an entire cell population. The observed upregulation of *MYC* in MCF7 TDP2KO cells could stem from either (1) a scenario in which only a subset of cells robustly upregulates *MYC,* creating a bimodal distribution that elevates the population average or (2) a situation where *MYC* is induced more robustly in the vast majority of MCF7 TDP2KO cell population than in MCF7 cell population, resulting in the normal distribution. The first scenario suggests that, for those DSBs that are involved in *MYC* regulation after E2 treatments, TOP2-induced DSBs are primarily re-ligated by TOP2 itself, and TDP2 plays a role in repair only in a subset of cells with abortive TOP2ccs. The second scenario implies that TDP2 is programmed in the repair of TOP2-induced DSBs that regulate *MYC* expression. These possibilities can be distinguished by single-cell RNA-seq (sc RNA-seq) experiments.

Four groups of cells were subject to scRNA-seq: MCF7 parental cells, MCF7 parental cells treated with E2 for 2 hours, MCF7 TDP2KO cells, and MCF7 TDP2KO cells treated with E2 for 2 hours. Quality control and normalization of scRNA-seq data were executed using the Seurat pipeline [30]. Following the exclusion of poor-quality cells and genes, the transcriptome data comprised 354 genes in E2-treated MCF7, 585 genes in untreated MCF7, 708 genes in E2-treated TDP2KO, and 399 genes in untreated TDP2KO. For the filtered and normalized data, we conducted two independent comparisons: E2-treated MCF7 versus untreated MCF7 and E2-treated TDP2KO versus untreated TDP2KO. Applying an adjusted p-value <0.05 and log fold change ≥ 0.1 (1.11 fold), we identified a total of 111 genes that were differentially expressed between untreated and E2-treated MCF7; 100 upregulated and 11 down-regulated genes, respectively (**Table S4**). For TDP2-KO cells, E2 treatment resulted in gene expression changes in 327 genes: 138 up- and 189 down-regulated genes (**Table S5**).

Consistent with the findings from digital PCR and bulk RNA-seq, *MYC* expression was induced by E2 treatment in both MCF7 TDP2KO and MCF7 cells, with log2-fold changes of 0.43 (1.54 fold) and 0.17 (1.19 fold), respectively (**Fig. 4A**). Notably, the *MYC* induction by E2 was more pronounced in MCF7 TDP2KO cells than in MCF7. Normalized expression data revealed that MCF7 TDP2KO cell populations displayed continuous distributions without distinct subpopulations and uniquely included cells with very high *MYC* expression (normalized expression >1.5) after E2 treatment (**Fig. 4B**). Histograms of raw read count were consistent with the skewed unimodal distribution across all four populations (**Fig. 4C**). The normal distribution of expression values at a single cell level suggests that most cells in the population contributed equally to the mean increase, providing supports for the scenario (2).

We next examined whether the enhanced transcriptional response of the *MYC* in MCF7 TDP2KO cells is associated with cellular processes, such as cell cycle phases. Cell cycle phases are considered confounding factors in single-cell studies [31]. To address this concern, we allocated individual cells to G1, S, and G2/M phases using the cell cycle prediction tool based on single transcriptome data [32]. We then examined whether cell cycle distribution might confound the observed difference in *MYC* transcriptional responses by collecting *MYC* expression profiles with and without E2 in each cell cycle phase (Fig. 4D). Based on the cell cycle phase prediction, we found that MCF7 TDP2KO cells exhibited modestly higher enrichment in G1 cell populations than MCF7 cells; >60% of TDP2 KO cells were in G1, while 50% of MCF7 cells were in G1 (**Fig. S2**). However, the difference of *MYC* transcriptional induction by E2 was less pronounced between MCF7 and MCF7 TDP2KO cells in G1 cells (1.26-fold with p<0.0001 in MCF7 vs. 1.47-fold with p<0.0001 in MCF7 TDP2KO, Wilcoxon rank-sum test) than in S (1.08-fold with p=0.0332 vs. 1.52-fold with p<0.0001) and G2M cells (1.24-fold with p=0.0014 vs. 1.88-fold with p<0.0001) (**Fig. 4C**). Therefore, the difference in the distribution of cell cycles was not a major contributor to increased *MYC* induction in MCF7 TDP2KO cells. Instead, cell cycle profiling revealed that the increased response to E2 would likely stem from cells in S/G2M phases.

### The dominance of upregulated genes by the short period of E2 treatment

The global transcriptional responses to E2 have been extensively studied in the MCF7 cell line, and both upregulated and downregulated genes have been documented. However, variations in the amounts of E2 and treatment durations across experiments, as well as the differences in genomic platforms (e.g., microarray, RNA-seq, and scRNA-seq), could influence the repertoire of E2-responsive genes, as previously discussed [33, 34]. To establish a reliable set of E2 (10nM)--responsive genes at an early time point (2 hours), we sought the intersections between different genomic platforms, bulk RNA-seq, and scRNA-seq.

Bulk RNA-seq analysis identified a large number of differentially expressed genes, including 2021 for MCF7 and 1034 for MCF7 TDP2KO, between untreated and E2-treated cells. In contrast, the numbers of differentially expressed genes in our scRNA-seq data were modest: 111 genes for MCF7 and 327 genes for MCF7 TDP2KO (**Fig. 5A**). Thirty-two genes were common between bulk and single-cell RNA-seq in MCF7, while 62 genes were shared in MCF7 TDP2KO cells. The majority of these common genes were upregulated genes in both cell lines (**Fig. 5B; Tables S6** and **S7**). In MCF7, 28 genes were upregulated, and four genes were repressed, with one gene, GPRC5A, in opposite directions between bulk- and sc-RNA-seq. In MCF7 TDP2KO, 58 genes were upregulated, and four genes were repressed. There were 8 common differentially expressed genes between MCF7 and MCF7 TDP2KO, including *MYC*, *CCND1*, *NRIP*, *SH3BP5*, *IGFBP4*, *RET*, *NBPF1* and *CA12* (**Fig, 5C**). Notably, *MYC* and *CCND1* are well-established oncogenes in breast cancer [35–38]. In summary, a reliable set of estrogen-responsive genes at an early time point (2h) were predominantly upregulated genes, with TDP2 potentially regulating responses to E2 for both *MYC* and *CCND1*.

**Figure 5.**
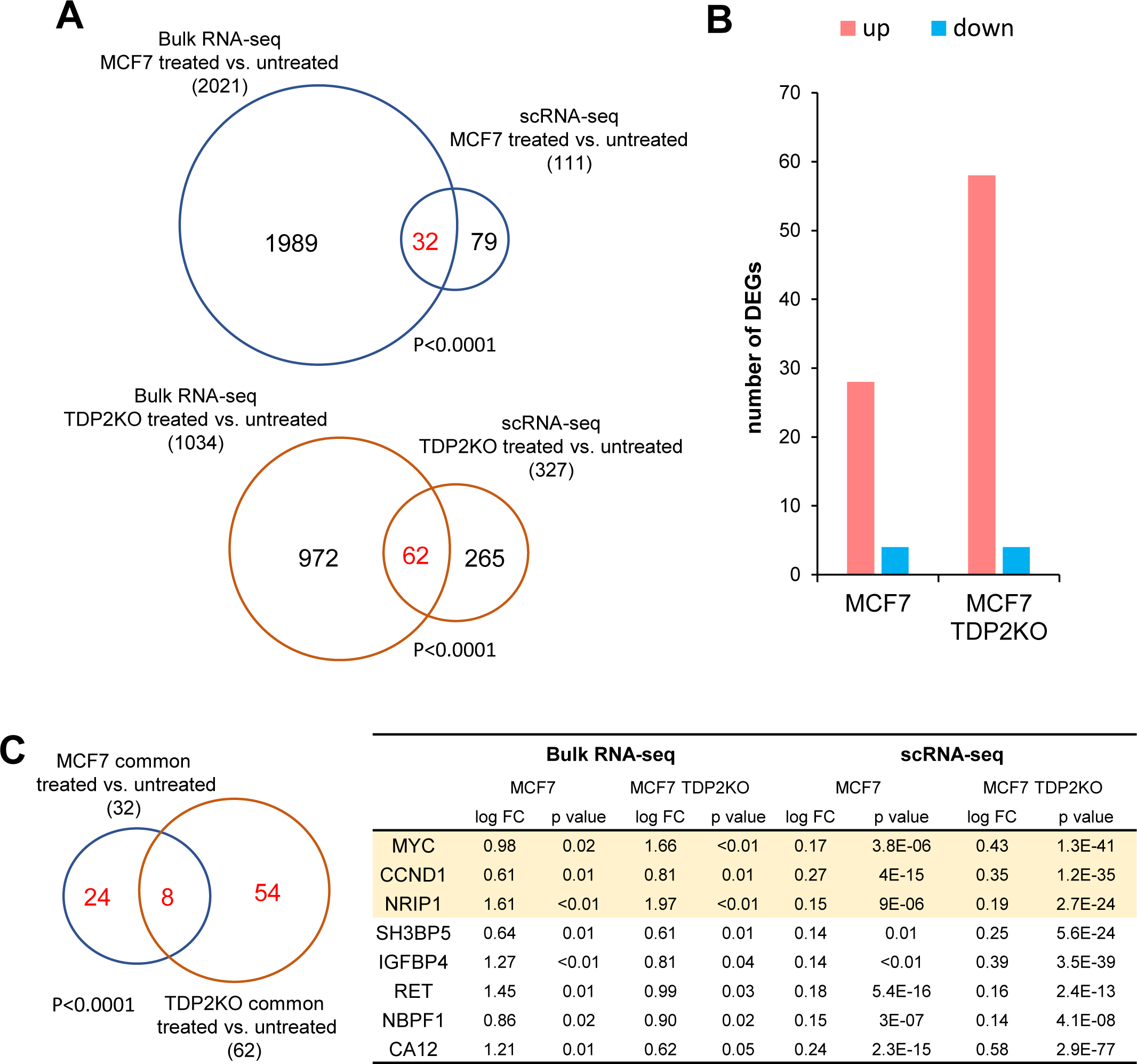
Commonly upregulated genes between bulk- and single-cell RNA-seq. (A) Venn diagrams showing the differentially expressed genes: The differentially expressed genes by E2 treatment were presented for MCF7 and MCF7 TDP2KO cells. 32 genes in MCF7 and 62 genes in MCF7 TDP2KO are common between the two platforms. Significances of overlap were assessed using Fisher’s exact test. (B) Bar plots showing the number of common differentially regulated genes s by E2 (2 hours): 28 and 4 genes were commonly up- and down-regulated in E2-treated MCF7 compared to untreated MCF7. Similarly, 58 and 4 genes were commonly up- and downregulated in E2 treated MCF7 TDP2KO compared to untreated MCF7 TDP2KO. (C) Venn-diagram and a Table of commonly differentially expressed genes: Depicting 32 and 62 commonly differentially regulated genes from bulk and single cell RNA-seq data, along with the table highlighting 8 overlaping genes. Significance of overlap was determined using Fisher’s exact test.

## Discussion

In this study, we assessed the transcriptional responses to 17beta-estradiol (E2), an abundant form of estrogen, both *in vitro* and *in vivo*, in ER-positive breast cancer cells and breast epithelial cells in mice. The transcriptional response to estrogen is known to involve topoisomerase 2 (TOP2) [3], which creates transient DNA double-strand breaks, relieves torsional stress, and reseal the ends. Recent studies have highlighted the roles of DNA repair proteins, including TDP2, Mre11, BRCA1, and NHEJ proteins, in the repair of TOP2-induced DSBs [15–18]. We aimed to gain insights into how these proteins contribute to the biological processes involving TOP2-induced DSBs. To do so, we monitored transcriptional response to E2 in TDP2 wild-type and TDP2KO MCF7 and T47D cell line. Defects in the repair of TOP2-induced DSB have been shown to upregulate transcriptional response to external stimuli [11]. Therefore, we anticipated that the extent of changes in the E2 response would be proportional to the fraction of the population experiencing repair defects due to the absence of TDP2.

We confirmed that transcription of an oncogene *MYC* responded to E2 (10 nM) very early, as early as 1 hour after the addition of E2 (**Fig. 1D**). The early response was inhibited when the catalytic activity of TOP2 was blocked by ICRF-193 in a dose-dependent manner in both MCF7 and T47D cell lines (**Fig. 1D**), strongly suggesting that TOP2 plays a crucial role in the early transcriptional response of *MYC*. The transcriptional response is most likely through TOP2-induced DSBs, as the TOP2 poison etoposide enhanced the transcriptional response (**Fig. 2E**). This notion was further supported by the *in vitro* experiments with cell lines deficient in TDP2 using various platforms for gene expression measurements (**Fig. 2, 3** and **4**). The enhanced *MYC* response was also observed in cells treated with an NHEJ inhibitor, indicating that unrepaired TOP2-induced DSBs were likely the cause of upregulation (**Fig. 2E**). Importantly, we detected the upregulation by the techniques that measured the average gene expression level in the entire cell population, such as digital PCR and bulk mRNA-seq. Therefore, TDP2 seems to be involved in the transcriptional regulation of *MYC* in a substantial fraction of the cell population.

Using scRNA-seq, we further revealed that the observed enhanced response of *MYC* in MCF7 TDP2KO cells, compared to MCF7 cells, was driven by the population-wide upregulation of *MYC* by E2. This was evidenced by the unimodal distribution of *MYC* mRNA in MCF7 TDP2KO cells (**Fig. 4C**). The unimodality of gene expression was seen for *MYC* and other commonly upregulated genes. In a previous study of scRNA-seq for the E2-stimulated transcriptional response in MCF7 cells, a large proportion of genes showed a bimodal distribution of gene expression [39]. However, the smaller numbers of cells captured for scRNA-seq in that study (87 MCF7 cells) compared to our study (354 cells from MCF7 with E2, and 585 cells from MCF7 without E2) may contribute to the difference. With these results along with the TOP2-dependent E2 response of *MYC* (**Fig. 1E**), we speculate that TDP2-dependent repair of TOP2-induced DSBs is programmed to regulate *MYC* expression when stimulated by E2. Unrepaired DSBs could potentially suppress the transcription of nearby genes [40–42]. Conversely, the accumulation of DNA damages at stimuli-induced genes would promote transcription by releasing PolII from pausing [11]. Also, unrepaired DSBs far from the target genes, such as enhancer elements, would not directly disrupt the elongation of target gene mRNA. This scenario could apply to *MYC,* whose enhancer elements are dispersed within a few Mb gene deserts surrounding *MYC* [43, 44]. Notably, an enhancer located at 135kb distal to *MYC* (Myc+135kb enhancer) has an ER binding site and responds to E2 treatments [45]. The estrogen-induced expression of enhancer RNA and the physical interaction between enhancer and *MYC* promoter was greatly promoted when the repair of TOP2-induced DSBs was inhibited by an ATM inhibitor (KU-55933). Although ATM and TDP2 function independently to repair etoposide-induced DSBs [46], and spatial and temporal regulations of these two repair pathways remain unclear, TDP2 may also regulate the estrogen-induced activation of *MYC* enhancers through the repair of TOP2-induced DSBs.

scRNA-seq also enabled us to investigate the responses to E2 within distinct cell cycle phases. We found that TDP2 KO cells responded to E2 more robustly in the S and G2/M phases than in the G1 phase. There are two pathways for the removal of abortive TOP2ccs from the ends: TDP2 and Mre11/BRCA1. It is currently unclear how these two pathways delineate their tasks in removal. BRCA1 supports the nuclease activity of Mre11 to remove abortive TOP2ccs [17], which was shown to be active in G1 cells. Considering the robust upregulation of *MYC* in S and G2/M phases in TDP2-KO cells, we speculate that these two pathways repair DSBs with abortive TOP2cc in different cell cycle phases. A recent study showed that TDP2 binds to K63 and K27 poly-ubiquitinated cellular proteins, and this binding stimulates TDP2-catalyzed removal of TOP2 from DNA [47]. The identity of the K27 and/or K63 poly-ubiquitinated proteins bound to TDP2 is not known. However, it is conceivable that such a protein could regulate TDP2 activity in a cell-cycle-dependent manner.

We utilized both bulk and scRNA-seq for evaluating the responses of E2-responsive genes at a 2-hour time point. The ER-positive MCF7 cell line has been employed to study E2-responsive genes [33, 34]. However, these studies varied in their experimental conditions, including measurement platforms, E2 concentrations, and treatment durations, resulting in substantial variations in the numbers and spectra of E2-responsive genes. Additionally, genetic variations among MCF7 sublines [48] may contribute to the variability in results. To address this, we employed the same clones and maintained identical experimental conditions across two RNA-seq platforms to minimize variabilities associated with the experimental design. The genes called as common genes between the two platforms were strongly restricted by scRNA-seq data, as only small fractions of differentially expressed genes detected by bulk RNA-seq were also called by scRNA-seq. This discrepancy is likely attributed to the inherent heterogeneity of scRNA-seq data, which often includes numerous zero values for a substantial fraction of genes [49]. Nevertheless, these common differentially expressed genes showed a pronounced bias toward upregulated genes. Notably, these genes included oncogenes *MYC* and *CCND1,* frequently upregulated in ER-positive breast tumors. Estrogen induces mammary ductal hyperplasia through MYC in mice deficient in NHEJ, which is downstream of TDP2-mediated repair of TOP2cc. CCND1 is more frequently upregulated in ER-positive tumors than other subtypes and promotes cell cycle progression [50, 51]. By participating in the regulation of these oncogenes, TDP2 would emerge as a potential suppressor for the most common subtype of breast tumors.

## Materials and Methods

### Cell culture and digital PCR

MCF7 and MCF7 TDP2KO cells were maintained with MEM with 10% FBS. T47D, T47D TDP2KO, and MDA-MB-231 cells were maintained in DMEM with 10% FBS. TDP2KO cells were created by targeting exon 2 of the *TDP2* gene (5’-GGCTCAGAGATGGTTTCAGGT-3’). The targeting vector was constructed using pSpCas9(BB)-2A-GFP(PX458), a gift from Feng Zhang (Addgene plasmid #48138).

Etoposide sensitivity was assessed through the colony formation assay. In brief, 3000 cells were plated in each well of a 6-well plate, and cells were cultured for 8 days with or without etoposide before staining with crystal violet. Experiments were conducted in triplicate. Colonies were counted using the Hybrid Cell Function of the Keyence BZ-X700 microscopy (Keyence).

RNA was extracted using the RNeasy mini kit, and first-strand cDNA synthesis was performed using Superscript III (Invitrogen). Cells were cultured in phenol red-free medium supplemented with 5% FBS-charcoal stripped for 24 hours then treated with 10 nM 17β-estradiol for either 1, 2 or 6 hours before RNA extraction. NU7441 was added to the media 3 hours before adding 10 nM E2. Etoposide was added to the media 1 hour before adding E2. Digital PCR was conducted using the QuantStudio 3D Digital PCR System. A TaqMan probe labeled with FAM (MYC-Hs00153408, IL20-Hs00218888 or OLFML3-Hs01113293) and VIC (TBP-Hs00427620 or TFRC-Hs00951083) were mixed in a reaction, and the amount of transcript (copies/microliter) was determined using the Analysis Suite (Thermo Fisher Scientific).

### Mouse experiments

*Tdp2 Wild-type* and *Tdp2^-/-^* strains originated from the C57BL6/J strain. E2 (300 μg/kg body weight) was administered via intraperitoneal injection (IP) into three independent two-month-old female mice at a volume of 100 μL. As a control, an equivalent volume of a solvent (PBS) is injected.

The mice were sacrificed to harvest mammary gland tissues and frozen blocks were prepared. After a brief wash of the isolated tissues with cold PBS, the tissues were fixed with paraformaldehyde (4%, Cat# 163-20145, Wako, Japan) for 15 min at 4° C and then washed three times with 1x PBS. To preserve tissue integrity, incubation was performed in 30% sucrose (in PBS) for 30 min to 3 hours. Tissues were then embedded in optimal cutting temperature (OCT) compound (Cat# 4583, Sakura, Japan) into Cryomolds (Cat# 4565, Sakura, Japan). For cryo-sectioning of frozen blocks, a cryostat (CM1850, Leica, Germany) was set at -25° C and sectioned into 10 μm slices. The slides were then heated at 55° C for 10 to 30 min to dry. Dried slides were washed three times with Tween-20 in PBS (0.05 %, PBS-T). For blocking, 1 % BSA and 5 % goat serum in PBS-T were used for 3 hours at room temperature, followed by washing once with PBS-T. Slides were then incubated overnight at 4° C with both α-Cytokeratin-8/18 (1/10, rat monoclonal, University of Iowa, US) and α-Myc antibody (1/200 [Y69], ab32072, rabbit monoclonal, Abcam). After washing with PBS-T, slides were incubated with both α-rabbit (Alexa Fluor 546) and α-rat (Alexa Fluor 488) secondary antibodies (Molecular probe, US). Myc and Cytokeratin-8/18 signals were detected using LEICA SP8.

### Data preprocessing and normalization of bulk RNA-seq data

MCF7 cells, MCF7 cells treated with 10 nM E2 for 2 hours, MCF7 TDP2KO cells, MCF7 TDP2KO cells treated with 10 nM E2 for 2 hours, and etoposide-treated MCF7 with or without 10 nM E2 were subject to RNA-seq at the Cedars-Sinai Genomics Core. Biological duplicates were obtained for all of the experimental groups. Our quality control pipeline employed Illumina standards to assess base call quality and truncated low-quality reads. The quality of sequencing data was evaluated using MultiQC software [52]. Sequence alignment and quantification were performed using the STAR-RSEM pipeline [53, 54]. Reads overlapping exons in the annotation of Genome Reference Consortium Human Build 38 (GRCh38) will be identified. Genes failing to achieve raw read counts of at least two across all the libraries were filtered out and excluded from downstream analysis. Batch effects were corrected by ComBat [55]. The trimmed mean of the M-values normalization method (TMM) [56] (version 1.6.1) was used for calculating normalized count data.

### Differential expression analysis of bulk RNA-seq data

For the two comparisons - 1) E2 treated versus untreated in MCF7 cells and 2) E2 treated versus untreated in MCF7 TDP2KO cells - we employed the integrated hypothesis testing method, as previously reported [57]. Briefly, for each gene, two P-values were computed by conducting a two-tailed Student’s T-test and a two-tailed median different test using the empirical distributions that were estimated by random permutations of the samples. We then combined two P-values into a single combined P-value using Stouffer’s method. Multiple testing correction was done by Storey’s correction method [58]. Finally, the differentially expressed genes were selected with the ones having a combined p-value < 0.05 and a fold-change ≥ 1.5. The Gene Set Enrichment Analysis (GSEA) was employed to assess the enrichment of gene sets [28].

### Single-cell RNA-seq data preprocessing and differential expression analysis

Single-cell RNA-seq was performed using the 10X Genomics scRNA-seq approach at the Cedars-Sinai Genomics Core. The Cell Ranger Single-Cell Software Suite was used for demultiplexing, barcode assignment, and quantification of unique molecular identifiers (UMIs). Reads were aligned to the hg38 reference genome using a pre-built annotation package obtained from the 10X Genomics website. All lanes per sample were processed using the ‘cellranger count’ function. The output from different lanes was then aggregated using ‘cellranger aggr’ with –normalise set to ‘none.’ Quality control and analysis were conducted using the Seurat packages in R (version 3.2.2) [30]. Poor-quality cells, defined by the number of detected genes less than 2,000, were filtered out. In addition, all cells with 11.89% (three standard deviations) or more of UMIs mapping to mitochondrial genes were considered non-viable or apoptotic and excluded from the analysis. Genes detected in at least three cells were retained for further analysis, resulting in a total of 19,043 genes for the analysis. To ensure no index swapping occurred, cells with unique barcodes that appeared in more than one sample (non-unique barcodes) as ‘NULL’ were excluded for further analysis. Gene expression values were normalized through log normalization using the “NormalizeData” function in Seurat. We performed two comparisons: 1) E2 treated versus untreated in MCF7 cells and 2) E2 treated versus untreated in MCF7 TDP2KO cells. The “FindMarkers” function in Seurat was used with filtered data to identify differentially expressed genes between cell groups. We called differentially expressed genes with adjusted p-value <0.05 and log fold change ≥ 0.1 (1.11 fold). The fold change threshold was determined at the 95 percentile value of the empirical null distribution of fold changes, which was generated through 1,000 times random permutation of the cells.

### Statistical Analysis

We conducted Principal Component Analysis (PCA) to visualize the samples and assessed the distribution of the samples based on their gene expression profile. We used MATLAB (v.9.0; Mathworks, Natick, MA, USA), R (v.3.5), and Python (v.3.7) for bioinformatic analysis.

## Acknowledgment

We thank Cedars-Sinai Cancer, Applied Genomics, Computation and Translational Core and Flow Core for technical support. This work is supported by Cedars-Sinai CTSI Clinical Scholar Award (to N.M.), NIHR01CA149385, Cedars-Sinai Cancer Biology Development Fund and Institutional Support (to H.T.), Leukemia & Lymphoma Society (to N.J.) and Margie and Robert E. Petersen Foundation, The Fashion Footwear Charitable Foundation of New York, Inc., Avon Foundation and Associates for Breast and Prostate Cancer Studies (to A.G.).

## Data availability

Bulk and single-cell RNA-seq data are available in Gene Expression Omnibus (accession # GSE225423).

## Author Contributions

Conceptualization, N.M., M.K., S.T., S.Y.Y., and H.T.; Data collection, N.M., N.J., R.A.M., J.A, R.S., and H.T.; Data analysis and interpretation, N.M., M.K., R.A.M., S.Y.Y., and H.T.; Materials, F.C-L., S.Y., X.C.; Writing the manuscript, N.M., M.K., S.T., S.Y.Y., and H.T.; Funding Acquisition, N.M., A.E.G., and H.T.

