## Supplementary material for "TDP 2 modulates the expression of estrogen-responsive oncogenes": supp figures

### Slide 1
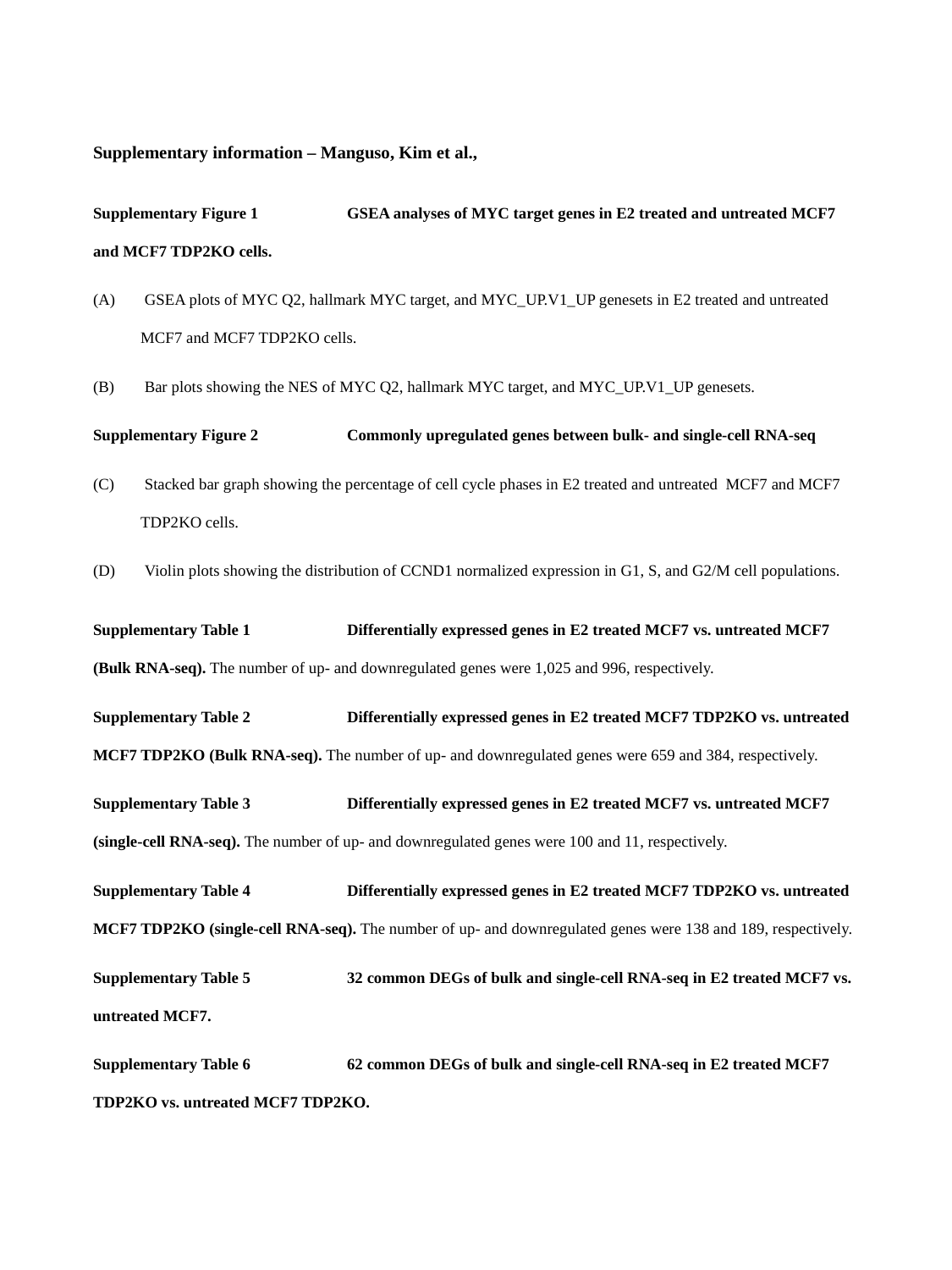

Supplementary information – Manguso, Kim et al.,
Supplementary Figure 1	GSEA analyses of MYC target genes in E2 treated and untreated MCF7 and MCF7 TDP2KO cells.
 GSEA plots of MYC Q2, hallmark MYC target, and MYC_UP.V1_UP genesets in E2 treated and untreated MCF7 and MCF7 TDP2KO cells.
 Bar plots showing the NES of MYC Q2, hallmark MYC target, and MYC_UP.V1_UP genesets.
Supplementary Figure 2	Commonly upregulated genes between bulk- and single-cell RNA-seq
 Stacked bar graph showing the percentage of cell cycle phases in E2 treated and untreated MCF7 and MCF7 TDP2KO cells.
 Violin plots showing the distribution of CCND1 normalized expression in G1, S, and G2/M cell populations.
Supplementary Table 1	Differentially expressed genes in E2 treated MCF7 vs. untreated MCF7 (Bulk RNA-seq). The number of up- and downregulated genes were 1,025 and 996, respectively.
Supplementary Table 2	Differentially expressed genes in E2 treated MCF7 TDP2KO vs. untreated MCF7 TDP2KO (Bulk RNA-seq). The number of up- and downregulated genes were 659 and 384, respectively.
Supplementary Table 3	Differentially expressed genes in E2 treated MCF7 vs. untreated MCF7 (single-cell RNA-seq). The number of up- and downregulated genes were 100 and 11, respectively.
Supplementary Table 4	Differentially expressed genes in E2 treated MCF7 TDP2KO vs. untreated MCF7 TDP2KO (single-cell RNA-seq). The number of up- and downregulated genes were 138 and 189, respectively.
Supplementary Table 5	32 common DEGs of bulk and single-cell RNA-seq in E2 treated MCF7 vs. untreated MCF7.
Supplementary Table 6	62 common DEGs of bulk and single-cell RNA-seq in E2 treated MCF7 TDP2KO vs. untreated MCF7 TDP2KO.

### Slide 2
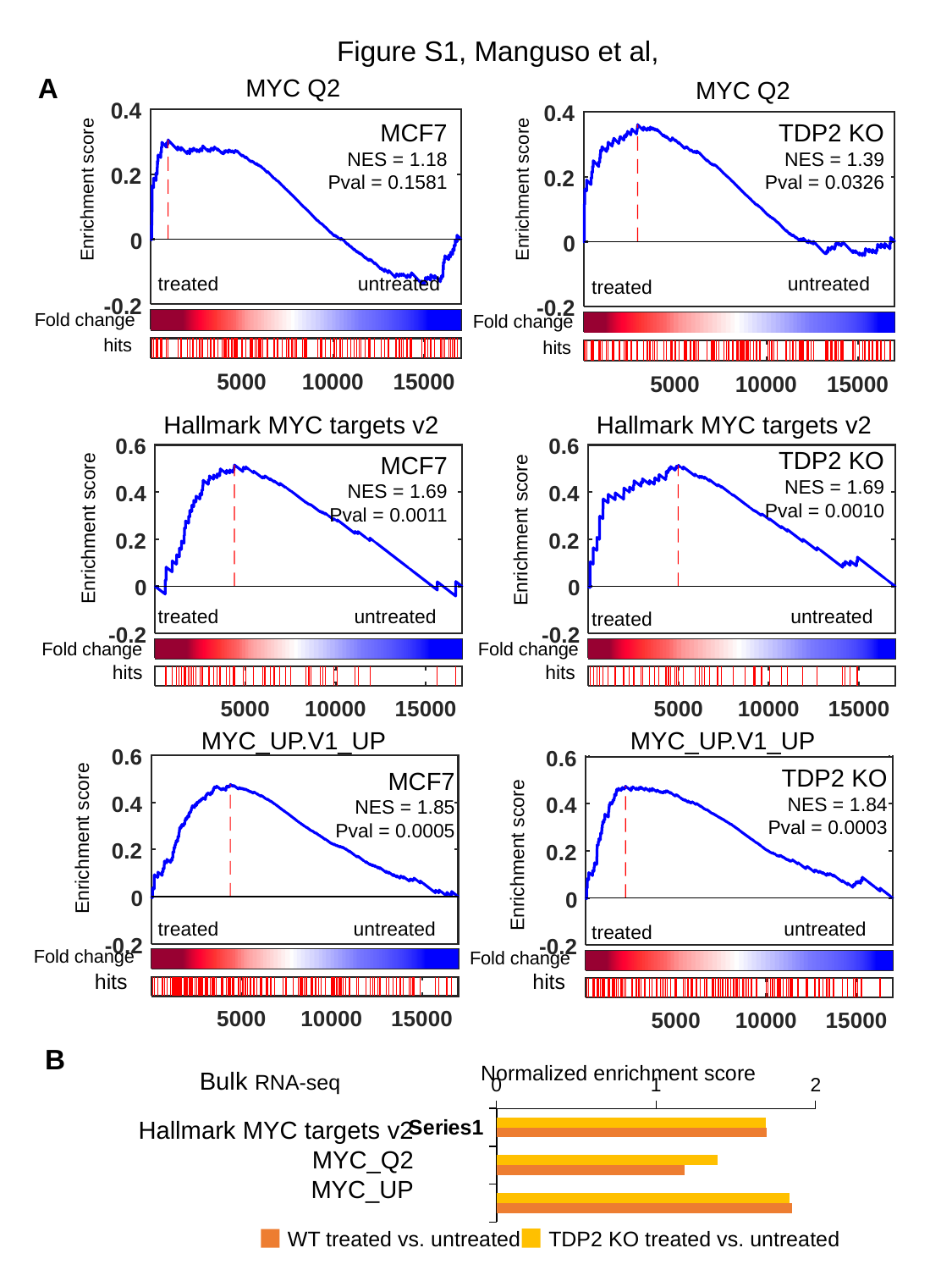

Figure S1, Manguso et al,
A
MYC Q2
MYC Q2
TDP2 KO
NES = 1.39
Pval = 0.0326
MCF7
NES = 1.18
Pval = 0.1581
Enrichment score
Enrichment score
treated
untreated
untreated
treated
Fold change
Fold change
hits
hits
Hallmark MYC targets v2
Hallmark MYC targets v2
TDP2 KO
NES = 1.69
Pval = 0.0010
MCF7
NES = 1.69
Pval = 0.0011
Enrichment score
Enrichment score
treated
untreated
untreated
treated
Fold change
Fold change
hits
hits
MYC_UP.V1_UP
MYC_UP.V1_UP
TDP2 KO
NES = 1.84
Pval = 0.0003
MCF7
NES = 1.85
Pval = 0.0005
Enrichment score
Enrichment score
treated
untreated
untreated
treated
Fold change
Fold change
hits
hits
B
Normalized enrichment score
Bulk RNA-seq
#### Chart
| Category | KO RNA-seq | WT RNA-seq |
|---|---|---|
| | 1.6874519545525306 | 1.6930982893706314 |
| | 1.3865016250419173 | 1.1804811286270513 |
| | 1.8389637787227155 | 1.8523203891158297 |Hallmark MYC targets v2
MYC_Q2
MYC_UP
WT treated vs. untreated
TDP2 KO treated vs. untreated

### Slide 3
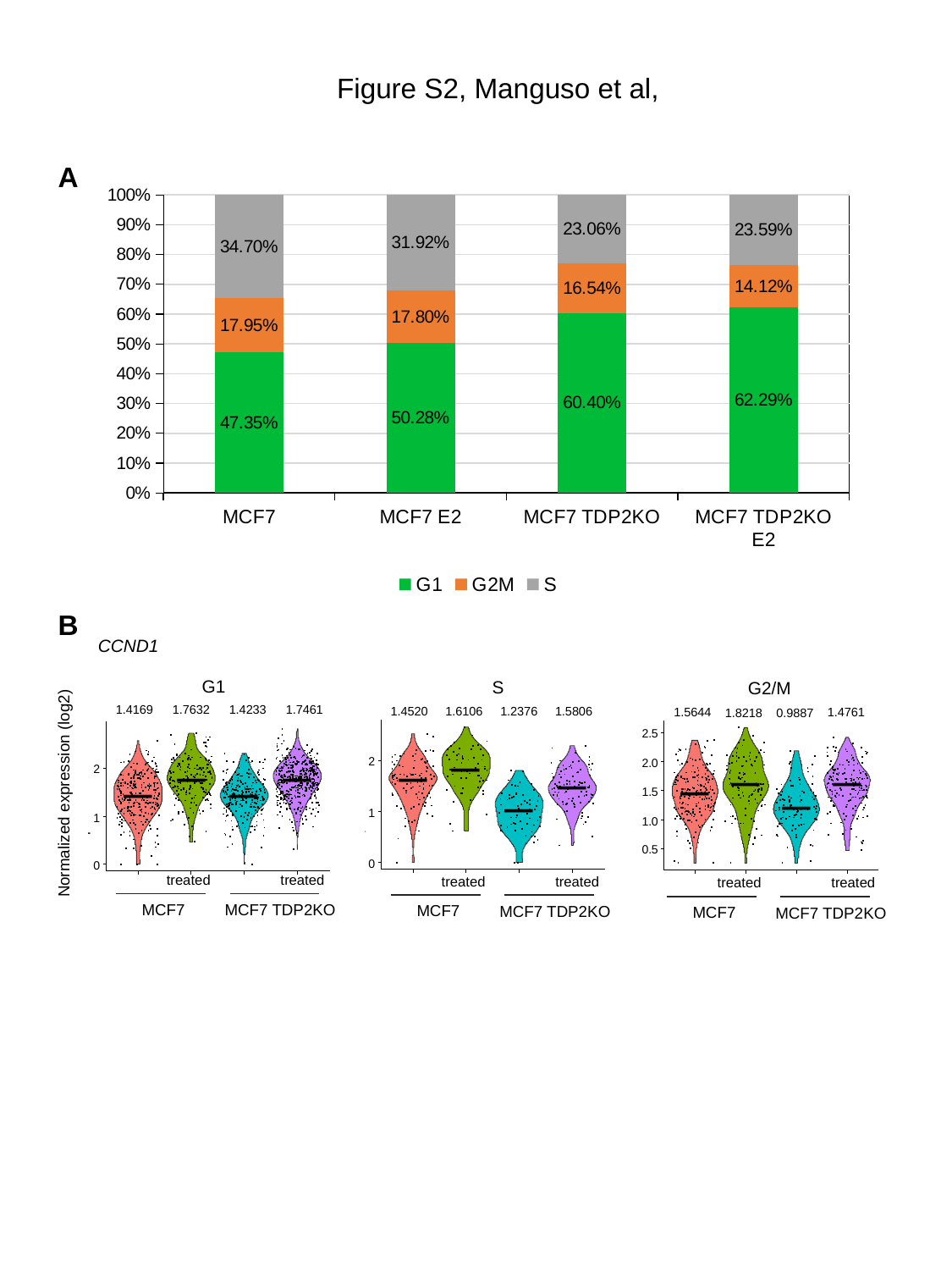

Figure S2, Manguso et al,
A
#### Chart
| Category | G1 | G2M | S |
|---|---|---|---|
| MCF7 | 0.4735042735042735 | 0.1794871794871795 | 0.347008547008547 |
| MCF7 E2 | 0.5028248587570622 | 0.17796610169491525 | 0.3192090395480226 |
| MCF7 TDP2KO | 0.6040100250626567 | 0.16541353383458646 | 0.23057644110275688 |
| MCF7 TDP2KO E2 | 0.6228813559322034 | 0.14124293785310735 | 0.23587570621468926 |B
CCND1
G1
S
G2/M
1.4169
1.7632
1.4233
1.7461
1.4520
1.6106
1.2376
1.5806
1.4761
1.5644
1.8218
0.9887
Normalized expression (log2)
treated
treated
treated
treated
treated
treated
MCF7
MCF7 TDP2KO
MCF7
MCF7 TDP2KO
MCF7
MCF7 TDP2KO
